# HRDex: a tool for deriving homologous recombination deficiency (HRD) scores from whole exome sequencing data

**DOI:** 10.1101/2022.09.08.506670

**Authors:** John Pluta, Ryan Hausler, Brad Wubbenhorst, Heena Desai, Susan M. Domchek, Katherine L. Nathanson, Kara N. Maxwell

**Author notes:** ***Correspondence to*** Kara N. Maxwell, MD PhD, Division of Hematology/Oncology, Department of Medicine, 421 Curie Blvd, Room 810, Philadelphia, PA 19104, USA.

## Abstract

**Background:** Breast and ovarian tumors in patients with biallelic *BRCA1* and *BRCA2* mutations either by germline mutations accompanied by allele-specific loss of heterozygosity (LOH) or truncal somatic mutations respond to PARP inhibition. The repair of double stranded DNA breaks in tumors these tumors leads to homologous recombination deficiency (HRD), which can be measured using a variety of genomic and transcriptomic signatures. However, the optimal biomarker for *BRCA* deficiency is unknown.

**Methods:** We developed HRDex to determine HRD and its composite scores from allele specific copy number data analysis of whole exome sequencing (WES) data and examined the discriminatory ability of HRDex and other genomic and transcriptomic measures to identify *BRCA* deficiency in breast and ovarian tumors from The Cancer Genome Atlas (TCGA).

**Results:** HRDex scores have high correlation with SNP array based HRD scores in both breast and ovarian cancers. HRDex scores have high discriminatory accuracy to distinguish *BRCA* deficient breast tumors, similar to SNP array based scores (AUC 0.87 vs 0.90); however, discriminatory ability for ovarian tumors was lower (AUC 0.79 vs 0.90). HRD-LST had the best discriminatory ability of the three composite HRD scores. HRDex had higher discriminatory ability for identification of *BRCA* deficiency than RNA expression based scores (eCARD, tp53, RPS and PARPi7) in breast and ovarian tumors. Tumor mutational burden (TMB) was associated with *BRCA* deficiency in breast but not ovarian cancer. Combining HRDex score with mutational signature 3 modestly increased discriminatory ability for *BRCA* deficient breast and ovarian tumors (breast: AUC 0.90 vs 0.87; ovarian: AUC 0.83 vs 0.79).

**Conclusions:** WES based HRD scores perform similarly to SNP array HRD scores, and better than other genomic or transcriptomic signatures, for identification of tumors with *BRCA* deficiency due to biallelic *BRCA* loss.

## Introduction

Tumors develop homologous recombination deficiency (HRD) due to a variety of mechanisms, including both germline and somatic inactivation of BRCA1 and BRCA2. Repair of double stranded DNA breaks in tumors with HRD occurs by error prone mechanisms leading to genomic instability which can be measured using a variety of genomic and transcriptomic signatures^1 2^. HRD has clinical relevance as when tumor cells are stressed with further DNA damage, for example due to stalled replication forks resulting from trapping of PARP on DNA by inhibitors, cell death occurs by synthetic lethality^3^. PARP inhibitors have FDA approved indications in ovarian, breast, pancreatic and prostate cancer patients with germline *BRCA1/2* mutations^4–11^. However, inactivation of a number of HR genes by genetic mutation or epigenetic silencing may lead to HRD in *BRCA1/2* wildtype tumors, but which specific HR genes should be interrogated is unclear^1^. On the other hand, not all *BRCA1/2* alterations may lead to HRD, ie in the absence of locus specific loss of heterozygosity^12,13^. Therefore, a more generalizable phenotypic readout of HRD is needed.

Biomarkers of *BRCA* deficiency have long been sought^14^. Array based comparative genomic hybridization (aCGH) signatures can distinguish tumors with copy number changes seen in germline *BRCA1* and/or *BRCA2* tumors^15,16^. Transcriptomic based measures of DNA repair deficiency have also been developed (ie eCARD^17^, PARPi7^18^, RPS^19^, tp53 score^2^). The most widely used HRD measure from tumors was originally developed using SNP arrays^20,21^. These HRD scores identify three discrete gross chromosomal abnormalities: genomic LOH (HRD-LOH)^22^, telomeric allelic imbalance (HRD-TAI)^23^, and large state transitions (HRD-LST)^24^. The combination HRD score has been extensively validated, demonstrating ability to predict *BRCA* deficiency^25^ and response to both platinum chemotherapy^26^ and PARP inhibitor^27,28^. However, at present, exome or other targeted based next generation sequencing (NGS) approaches, without aCGH, SNP arrays, or RNA expression analyses, are commonly performed clinically. Therefore, we developed an algorithm called HRDex to determine HRD scores from allele specific copy number data analysis of WES data.

## Methods

TCGA breast and ovary level 1 whole exome sequencing data (including BAM) were obtained from The Cancer Genome Atlas (TCGA) with a dbGAP approved project^29^. Downloaded TCGA BAM files were aligned to the hg19 assembly of the human genome. All exonic single nucleotide and insertion/deletion variants were identified using a combination of GATK^30^, Mutect^31^ and VarScan2^32^. Variants underwent initial quality control filtering according to GATK best practices^30^. Germline and somatic *BRCA1/2* alterations were identified as previously described^29^. *BRCA* deficient samples were defined as tumor-normal pairs with a germline loss of function *BRCA1* or *BRCA2* mutation with loss of heterozygosity (LOH) or a somatic *BRCA1* or *BRCA2* mutation with allele frequency (AF)>50%. Tumors with germline or somatic mutations in other genes in the homologous recombination pathway were excluded. This resulted in splitting of 708 TCGA breast cancer samples into 699 *BRCA* proficient and 39 *BRCA* deficient and splitting of 152 TCGA ovarian cancer samples into 99 *BRCA* proficient and 53 *BRCA* deficient.

HRDex was created to use an input file of genomic segments; where each segment is defined by its start and end position in physical coordinates with the copy number of each allele at said positions. Sequenza^33^ (v2.1.2, release date 10/10/2015) was used to create input files for the TCGA cohorts in this study. As the Sequenza segmentation algorithm can return false breaks (e.g. reporting two segments when in truth there is only one), if two adjacent segments had the same copy number and number of A and B alleles, and the gap between them was less than 3 Mb, this was considered a false break, and the two segments were joined. The length of the newly joined segment was defined as the distance between the start point of the first segment and the end point of the second.

To create the HRDex component scores and the sum, loss of heterozygosity (LOH) events were defined as segments that have a B allele copy number of 0, length greater than 15 Mb, and length less than 90% of the total chromosomal length. Large state transition (LST) events were defined as two adjacent segments with a gap of at least 3 Mb, where each segment is greater than 10 Mb and does not cross the centromere. Non-telomeric allelic imbalance (NTAI) events were defined as any segment that is not in either telomere, that does not cross the centromere, and has a segment size of greater than 11 Mb; segments with copy number equal to ploidy value were removed.

The ploidy of a tumor can be expressed as the number of genomic doublings that occur: p=2^n^; where *p* is the ploidy, and *n* is the number of genomic doublings. HRDex component scores were normalized to ploidy by dividing the total number of events by the number of genomic doublings. Ploidy was estimated from the data using Sequenza and averaged over the genome, therefore, a correction factor was included to account for continuous values. We defined a ploidy correction factor, *k*:

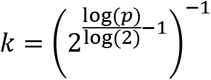

HRDex.LST and HRDex.LOH were normalized by multiplying the number of events by *k* (HRDex.LSTr and HRDex.LOHr). HRD.NTAI was normalized by subtracting the main copy number segments first, and then multiplying by *k* (HRDex.NTAIm). HRDex is the sum of the three component scores.

For public data analyses and comparisons of HRDex scores to SNP array based scores, SNP array based HRD scores, tumor mutational burden, aneuploidy score, expression signatures and mutational signatures for TCGA BRCA and OV datasets were obtained from Ref^2^.

ROC curves comparing the two data sets were computed, using leave-one-out cross validation to estimate out of sample performance. Area under the curves (AUC) of the two scores were statistically compared using deLong’s test.

## Results

HRD scores have been defined as a sum of three genomic events, LOH, LST and NTAI^25^. Because these events may increase in number based on increasing ploidy status of a tumor and not because of deficiency in the HR pathway, we tested whether there was a relationship between SNP array based HRD scores and ploidy in TCGA pancancer data^2^. The relationship between ploidy and SNP array based HRD scores differed between cancer types, with stronger correlations seen in certain cancer types such as uterine (UCEC), sarcoma (SARC), breast (BRCA) and stomach (STAD). Furthermore, certain cancer types had no correlation between HRD score and ploidy (**Supplementary Figure 1a**).

Because we noted a relationship between HRD and ploidy in certain cancer types, including breast, we created a tool to calculate ploidy-normalized HRD scores from allele specific copy number based analysis of whole exome sequencing data, called HRDex. HRDex is a freely available R package (https://github.com/maxwell-lab/HRDex), and outputs three component scores HRD.LOH, HRD.NTAI, HRD.LST and the sum of these three metrics, HRD.Score (**Supplementary Table 1**). HRDex scores normalized for ploidy remove the coorelation between HRD and ploidy, as shown for breast (**Supplementary Figure 1b**).

In order to test the HRDex algorithm, TCGA breast and ovary tumors were stratified into *BRCA* deficient and proficient tumors based on analysis of germline and somatic *BRCA1/2* mutations. Only tumors with germline or somatic *BRCA1/2* mutations with biallelic loss were considered “*BRCA* deficient”. Tumors with other germline or somatic mutations in the homologous recombination pathway, germline *BRCA1/2* tumors without biallelic loss, and tumors with low VAF somatic *BRCA1/2* mutations were excluded. Remaining tumors were “*BRCA* proficient”. HRD scores based on SNP-array data were an excellent classifier of *BRCA* deficiency in TCGA-Breast, as expected (AUC=0.9, **Figure 1a**). HRDex had a high correlation with SNP array based HRD and its component scores in TCGA breast tumors (r^2^ = 0.68-0.87, **Supplementary Figure 2a**). HRDex, before and after ploidy normalization performed almost as well as SNP array based HRD in TCGA-Breast, with AUCs of 0.87 and 0.88, respectively (**Figure 1a**). The performance of SNP array based HRD versus HRDex was not significantly different (*p* = 0.68). HRD.score and each component were significantly higher in *BRCA* deficient versus proficient breast tumors, with the most significant difference seen for HRD.LST (**Figure 1b**). Accordingly, the AUC for HRD.LST was similar to summed HRD.Score, although all component scores had AUC>0.80 (**Figure 1c**).

**Figure 1:**
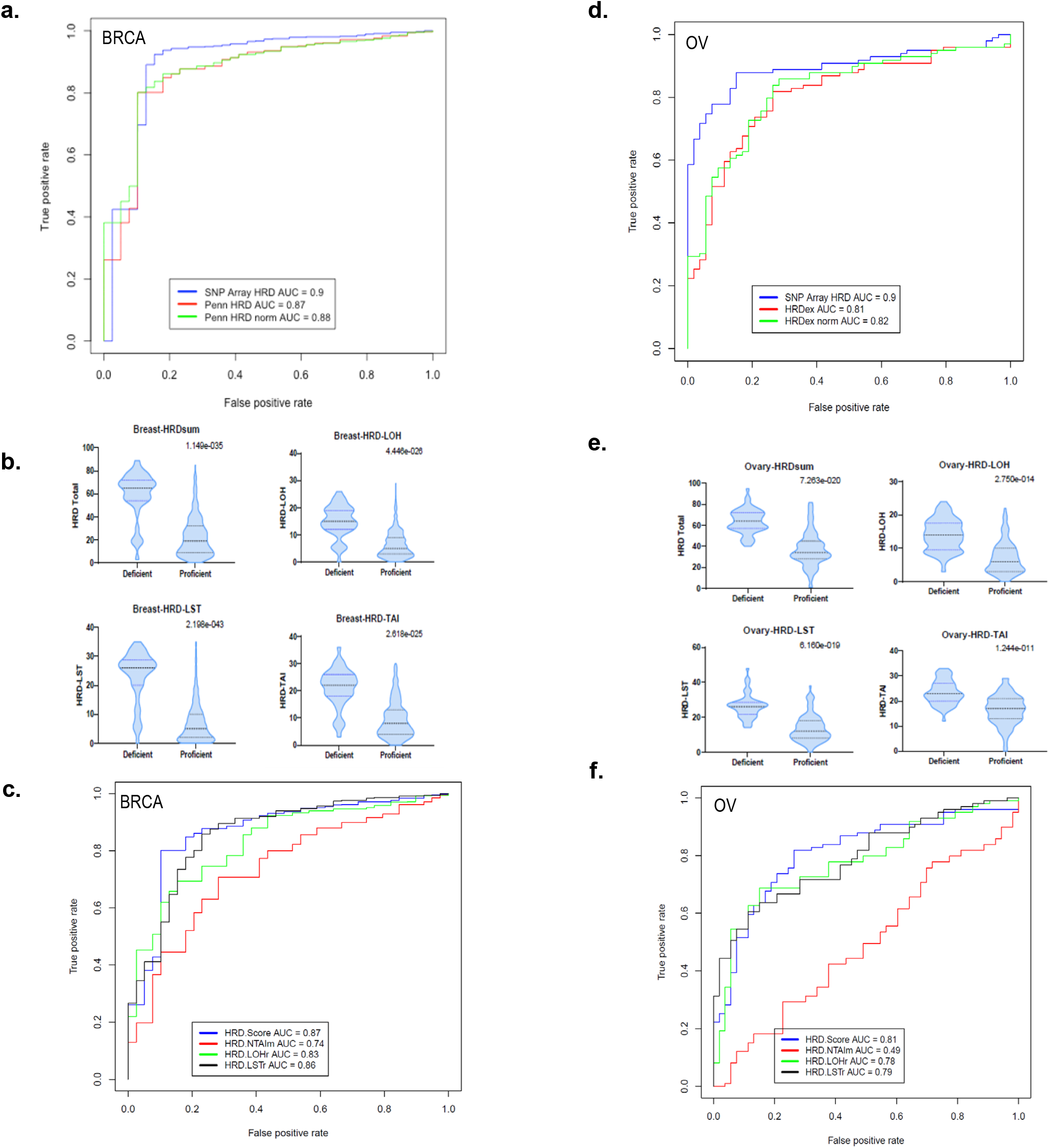
Performance of HRDex in TCGA breast and ovarian tumors. **(a)** Discriminatory performance of HRDex scores for *BRCA* deficient versus proficient TCGA breast tumors by HRDex, ploidy normalized HRDex, and SNP array based HRD. **(b)** Violin plots of component and total HRDex scores in *BRCA* proficient and deficient TCGA breast tumors. **(c)** Discriminatory performance of HRDex-total, HRDex-LOH, HRDex-LST, HRDex-NTAI component scores for *BRCA* deficient versus proficient breast tumors. **(d)** Discriminatory performance of HRDex scores for *BRCA* deficient versus proficient TCGA ovarian tumors by HRDex, ploidy normalized HRDex, and SNP array based HRD. **(e)** Violin plots of component and total HRDex scores in *BRCA* proficient and deficient TCGA ovarian tumors. **(f)** Discriminatory performance of HRDex-total, HRDex-LOH, HRDex-LST, HRDex-NTAI component scores for *BRCA* deficient versus proficient ovarian tumors.

HRD scores based on SNP-array were also an excellent classifier of *BRCA* deficiency in TCGA-Ovary, as expected (AUC=0.9, **Figure 1c**). HRDex had a high correlation with SNP array based HRD and its component scores in TCGA ovarian tumors, although lower than for breast (r^2^ = 0.42-0.79, **Supplementary Figure 2b**). HRDex, before and after ploidy normalization performed well in TCGA-Ovary, with AUCs of 0.81 and 0.82, respectively (**Figure 1c**). The performance of SNP array versus HRDex was not significantly different (*p* = 0.99). HRD.score and each component were significantly higher in *BRCA* deficient versus proficient ovary tumors, although the magnitude of differences were smaller in ovary compared to breast (**Figure 1e**). The AUC for the component HRD scores were lower in ovary, particularly for HRD-NTAI (**Figure 1f**).

Finally, we determined the discriminatory accuracy of the published DNA Damage Response (DDR) resource scores^2^ for distinguishing *BRCA* deficient versus proficient breast and ovarian tumors and compared to HRDex (**Table 1**). In TCGA-Breast, tumor mutational burden (TMB) had the highest AUC (0.98) for identifying *BRCA* deficient tumors; however, in TCGA-Ovary, TMB was a poor classifier (AUC=0.67). Aneuploidy score (defined as the sum of all aneuploid chromosome arms) was a poor classifier of *BRCA* deficient tumors in both TCGA-Breast and TCGA-ovary. Of DDR expression based measures, eCARD performed the best in TCGA-Breast (AUC:0.72); whereas PARPi7 performed the best in TCGA-Ovary (AUC:0.65). However, all DDR expression signatures had lower AUCs than genomic measures. In both TCGA-Breast and Ovarian cancer, mutational signature 3 was associated with *BRCA* deficiency (AUC 0.76 and 0.75). Combining HRDex or HRDex.LST score with mutational signature 3 modestly increased discriminatory ability for *BRCA* deficient tumors (breast: AUC 0.90 vs 0.87; ovarian: AUC 0.83 vs 0.79) versus the HRDex score alone.

**Table 1:**
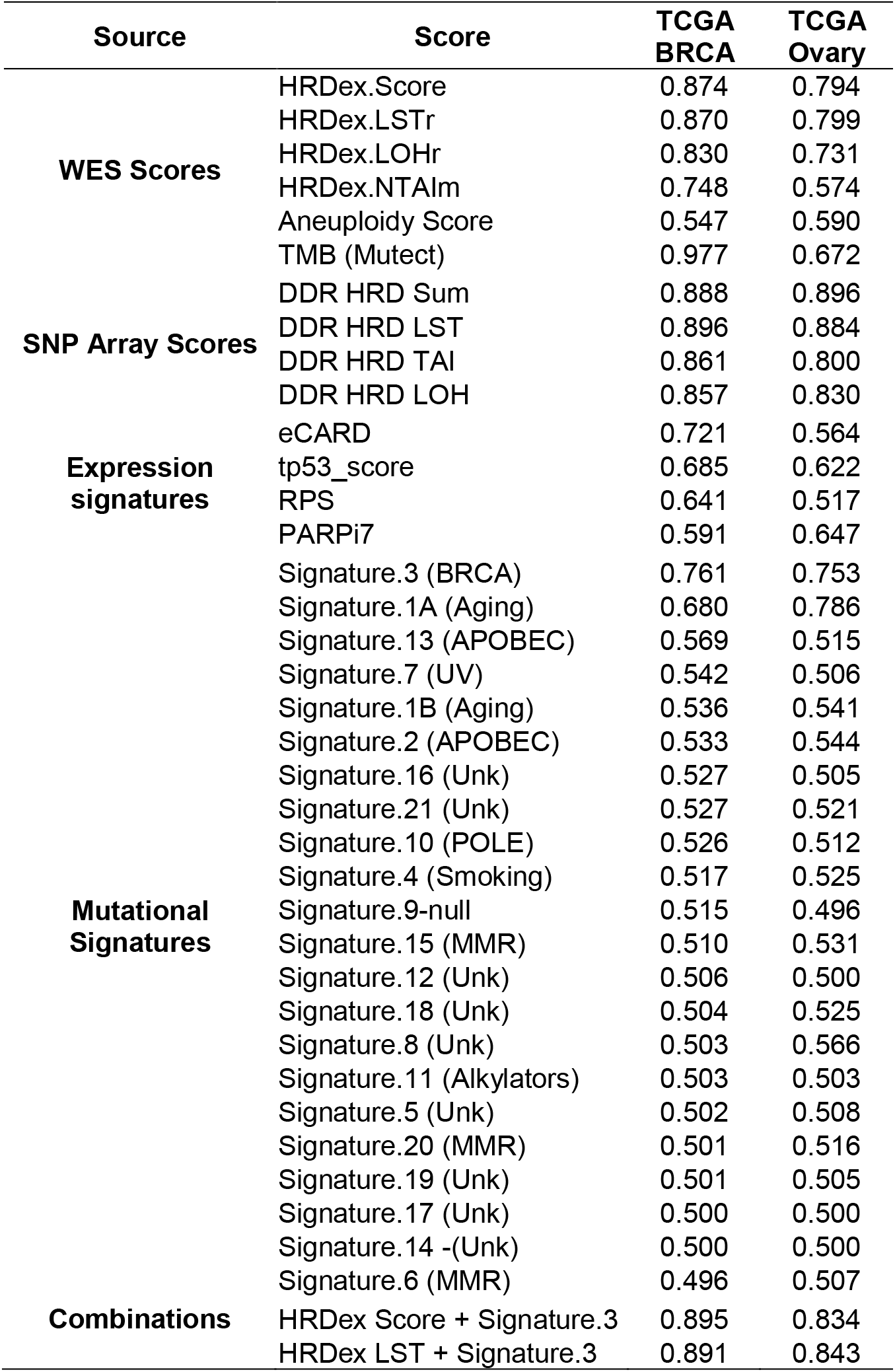
Performance of DDR scores in discriminating *BRCA* proficient and deficient tumors.

## Discussion

Homologous recombination deficiency (HRD) scores were originally developed using SNP array based data for the purpose of distinguishing germline *BRCA1/2* related tumors^22–24^. SNP arrays are rarely performed clinically or in genomics research due to the decreasing costs of next generation sequencing (NGS) based approaches. While whole genome sequencing (WGS) approaches are known to identify tumors with HRD^34–36^, WGS is still cost-prohibitive in most clinical and research settings. Given the large volume of whole exome sequencing (WES) data that is publicly available, we developed HRDex to derive HRD scores from allele specific copy number data derived from WES. We show that HRDex has comparable discriminatory accuracy to SNP array based HRD scores.

Currently, germline and/or somatic mutational analysis by NGS is the most commonly used tool to identify potential patients for platinum and/or PARP inhibitor eligibility^37^; however pan-cancer studies suggest mutational analyses alone will miss 30-40% of HRD tumors^34,38^. A number of trials have shown that HRD score is associated with response to neoadjuvant platinum based chemotherapy in breast cancer^20,25,26,39^. HRD scores may also predict responsiveness to platinum and PARP inhibitor in ovarian cancer^27,40^ and in tumors beyond breast and ovarian cancer^41^. Therefore, using NGS based HRD measures, such as HRDex or other NGS based approaches^42^, may expand identification of patients responsive to platinum or PARP inhibitors beyond just mutational analysis^37^.

It is important to note that the HRD score has not predicted platinum or PARPi responsiveness in all trials^43,44^, possibly due to reversion of HR function despite maintenance of genomic scars of prior HRD. Functional assays, such as RAD51 foci^45–48^ or ex vivo approaches^49^, may be superior in determining platinum and PARPi sensitivity but are not yet scalable in a clinical environment. A recent analysis suggests that incorporating *BRCA1* and *RAD51C* methylation assays into genomic analyses of all HR-related genes will find the vast majority of HRD tumors^50^; however, these assays are also not yet standardized.

HRD scores are important biomarkers of PARP inhibitor and platinum responsiveness, and useful in understanding the underlying biology of human tumors. HRD scores, such as HRDex, derived from readily available NGS based approaches, are useful in both clinical and research settings for the determination of HRD and study of PARP inhibitor responsiveness.

## Availability of data and material

HRDex scripts are available at: https://github.com/maxwell-lab/HRDex

## Acknowledgements

This work is supported by the National Cancer Institute (K08CA215312, KNM), the Burroughs Wellcome Foundation (#1017184, KNM), Basser Center for *BRCA* (KNM, SMD, KLN), V Foundation for Cancer Research (KLN), Gray Foundation (SMD, KLN) the Konner Family Foundation (KNM), and the Breast Cancer Research Foundation (SMD, KLN).

## Authors’ contributions

Study concept and design were performed by KNM and KLN. Data acquisition was performed by BW, and computational program created by BW, JP, and KNM. Data analysis and interpretation was performed by JP, RH, HD, KNM, SMD and KLN. Writing was performed by all authors.

---

**Supplementary Figure 1:**
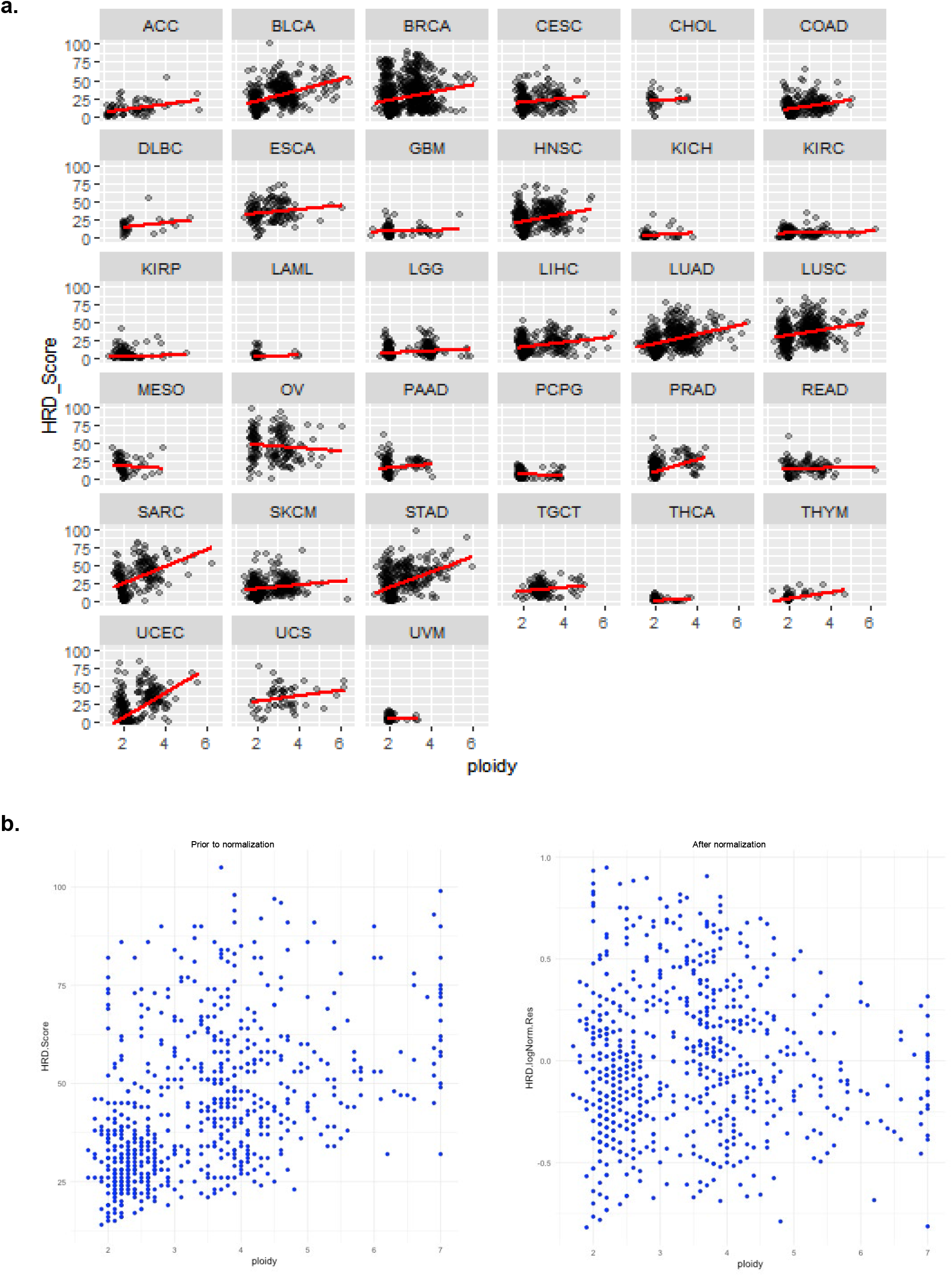
Development and performance of HRDex. (a): Analysis of pan-cancer TCGA data showing relationship of ploidy to raw HRD score as generated by SNP array data in different tumor types. (b) Correlation of HRDex score with ploidy prior to and after normalization in TCGA breast.

**Supplementary Figure 2:**
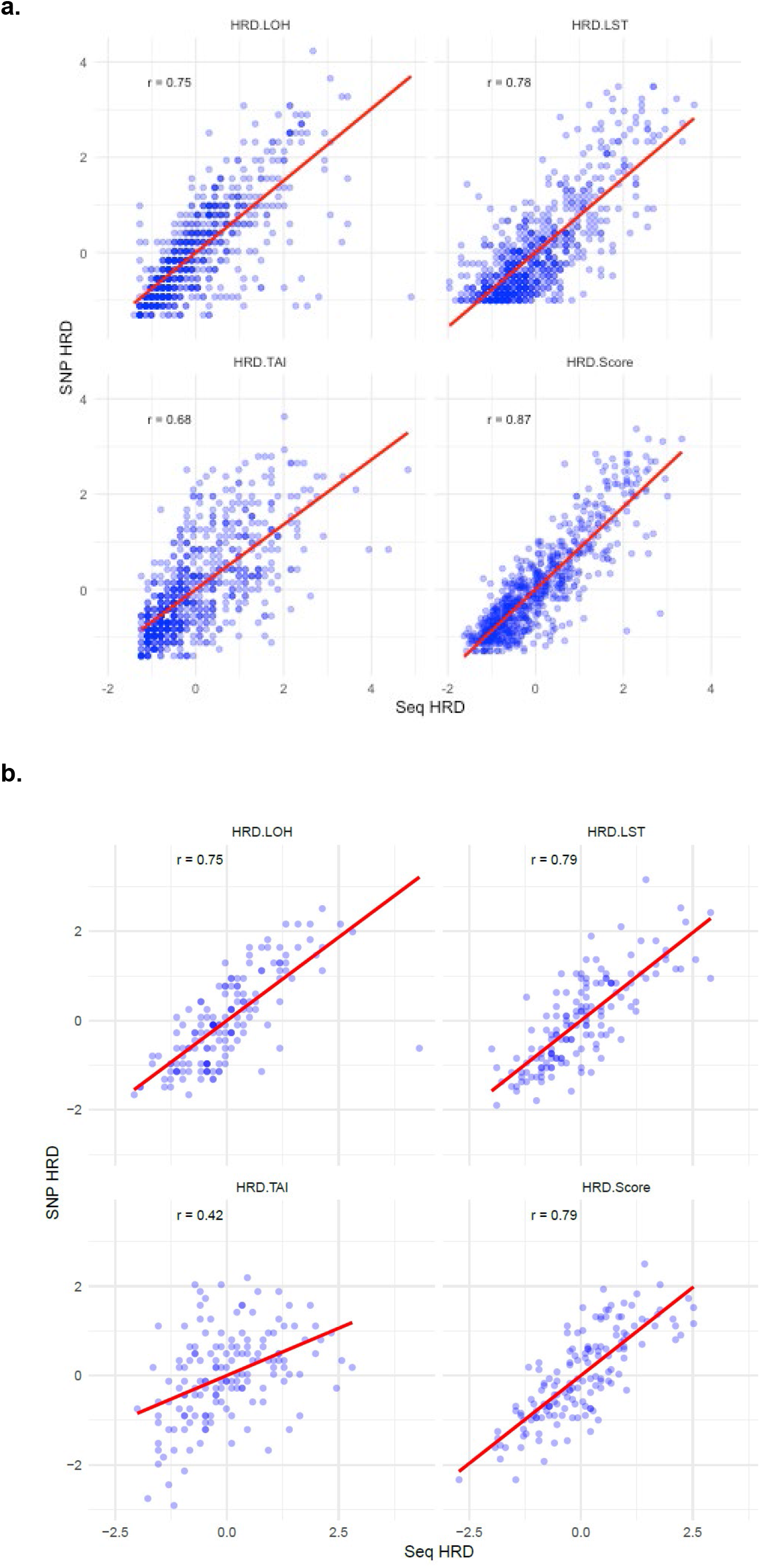
(a) Correlation plot of HRD score, LOH, LST and NTAI between pancancer SNP array data and HRDex derived for breast. (b) Correlation plot of HRD score, LOH, LST and NTAI between pancancer SNP array and HRDex derived for ovary.

**Supplementary Figure 3:**
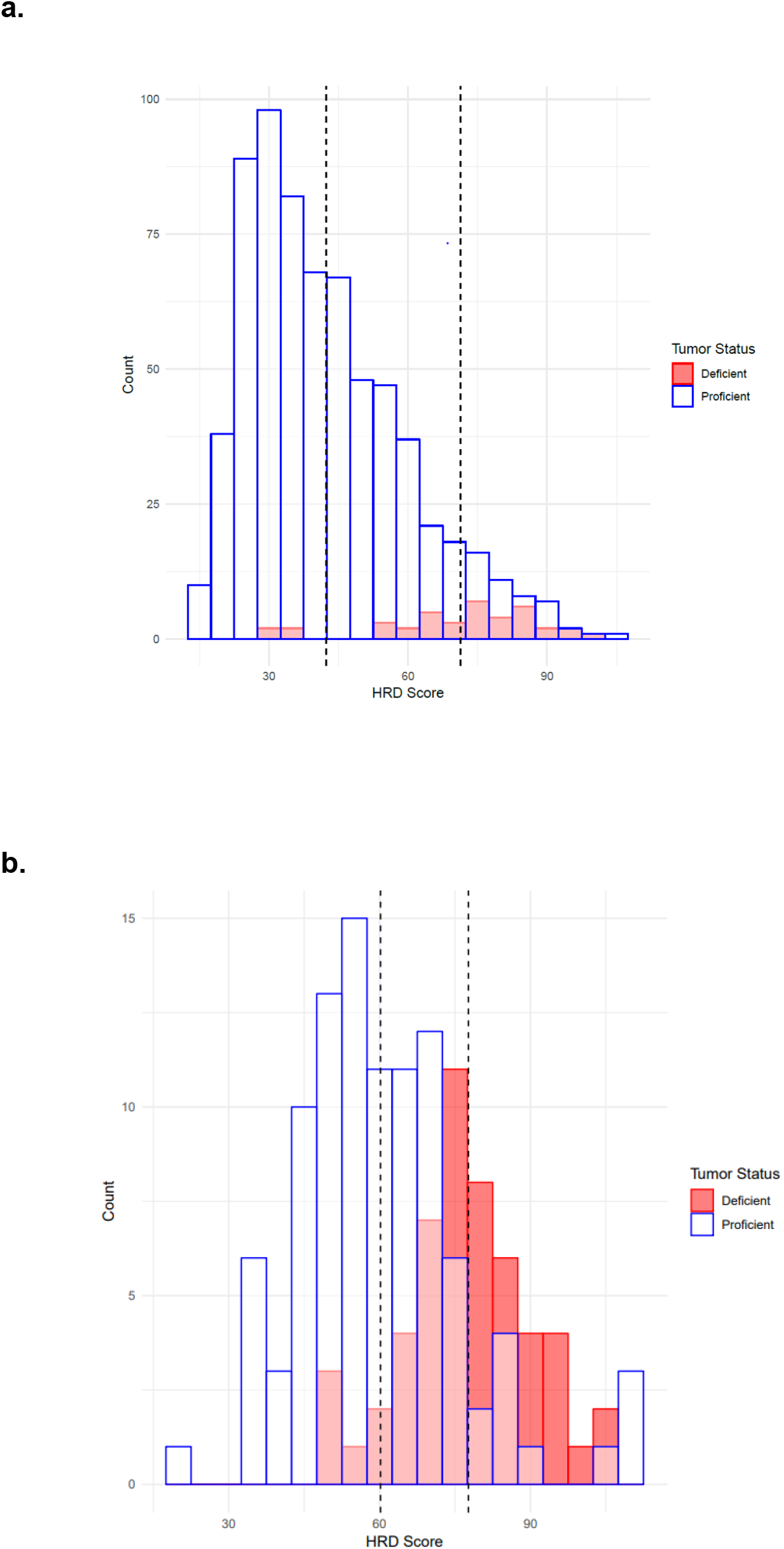
(a) Histogram of HRD scores by HRDex in TCGA breast (b) Histogram of HRD scores by HRDex in TCGA ovary

**Supplementary Figure 4:**
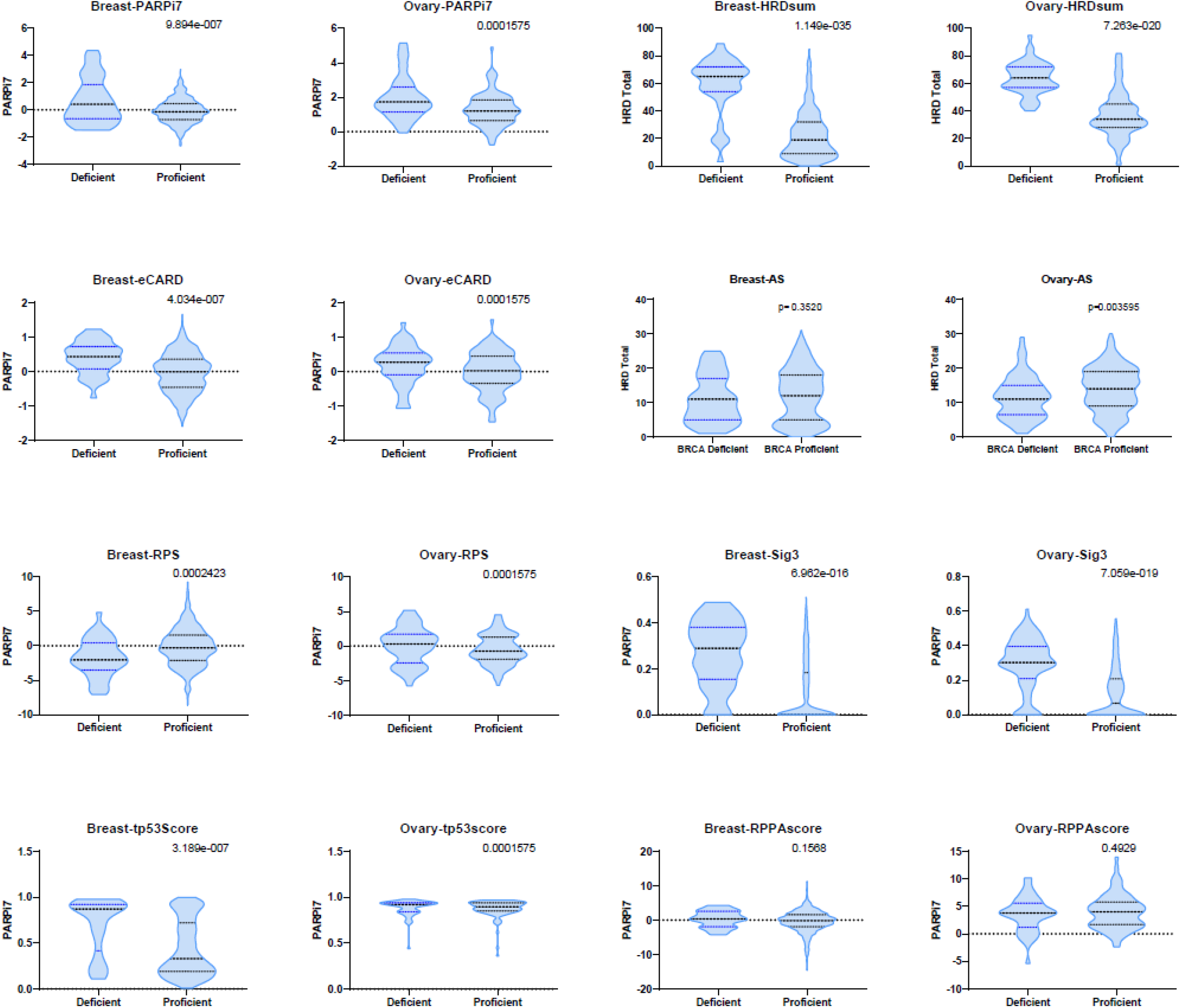
Violin plots of DNA Damage Response (DDR) scores from pan-cancer analysis in breast and ovarian tumors classified by *BRCA* status.

